# Five years of citizen science and standardized field surveys reveal a threatened urban Eden for wild bees in Brussels, Belgium

**DOI:** 10.1101/2021.03.10.434823

**Authors:** Nicolas J. Vereecken, Timothy Weekers, Leon Marshall, Jens D’Haeseleer, Maarten Cuypers, Pieter Vanormelingen, Alain Pauly, Bernard Pasau, Nicolas Leclercq, Alain Tshibungu, Jean-Marc Molenberg, Stéphane De Greef

**Affiliations:** Agroecology Lab, Université Libre de Bruxelles, Brussels, Belgium; Natuurpunt Studie, Mechelen, Belgium; Directorate Taxonomy & Phylogeny, Royal Belgian Institute of Natural Sciences, Brussels, Belgium; Natagora Bruxelles, Mundo-B, Bruxelles, Belgium

**Keywords:** Urban planning, urban green spaces, biodiversity, wastelands, vacant lands, brownfields, pollinators, wild bees

## Abstract

1. Urbanisation is often put forward as an important driver of biodiversity loss, including for pollinators such as wild bees. However, recent evidence shows that the mosaics of urban green spaces, and in particular certain categories of informal urban green spaces (IGS), can play an important role to help native wild bees thrive in cities.
2. Here, we describe the results of five years of citizen science and standardised field surveys of wild bees conducted at the Friche Josaphat, a 24-ha urban wasteland in the Brussels-Capital Region (Belgium). These field surveys were initiated following the planned restructuring and partial destruction of this site by the regional authorities.
3. We recorded a total of 2,507 specimens belonging to 127 species of wild bees, i.e. 60.5% of the 210 species recorded regionally, including nine that are threatened with extinction at national or European scales. The Friche Josaphat encompasses a significant share of the functional and phylogenetic diversity of wild bees known from the Brussels-Capital Region and is one of the most species-rich localities known to date for wild bees in Belgium.
4. Collectively, our results highlight the strong complementarity of citizen science and academic approaches in biodiversity surveys, and they reaffirm that wastelands are essential components of urban biodiversity. Our study stresses the need to provide biodiverse IGS with a formal status within the mosaic of urban green spaces, but also to acknowledge and safeguard their natural capital and the multiple ecosystem services they provide.

## Introduction

Urbanisation as a spatial process is *a priori* expected to have deleterious impacts on biodiversity, through its contribution to habitat fragmentation and the irreversible conversion of green spaces into impervious surfaces (e.g., McDonald *et al*., 2008; Vimal *et al*., 2012). Yet, parallel evidence suggests that some groups of organisms might actually thrive in cities (Miller & Hobbs, 2002; Araujo, 2003; Kühn & Klotz, 2006). By and large, our cities and megapolises are still home to relatively high numbers of native and sometimes rare or threatened species (e.g. Aronson *et al*., 2014), and also many exotic taxa (Fitch *et al*., 2019; de Souza e Silva *et al*., 2021; Taggar *et al*., 2021). These urban ecological networks are key to ecosystem function and resilience in cities under global change scenarios, but they also actively support the mental health, physical well-being and social interactions of present-day populations of urban dwellers (Barton & Grant, 2013; Bratman *et al*., 2019). Indeed, the benefits of regular access to (large) urban green spaces (UGS) were particularly exacerbated across the world urban centres during the recent COVID-19 pandemic and its associated travel restrictions (Pfefferbaum & North, 2020; Xie *et al*., 2020; Ahmadpoor & Shahab, 2021). For all these reasons, urban areas have become increasingly recognized as important targets for wildlife conservation (Goddard *et al*., 2010; Dearborn & Kark, 2010; Kowarik, 2011; Shwartz *et al*., 2014), as well as to comply with the UN Sustainable Development Goals aiming “to make cities and human settlements inclusive, safe, resilient, and sustainable” (United Nations, 2015; Apfelbeck *et al*., 2020).

The added value of cities for biodiversity conservation lies in their mosaics of typologically diverse urban green spaces (UGS), from playing fields to highly manicured environments such as managed forests, parks or cemeteries, to semi-natural landscapes, including urban nature reserves (Lepczyk *et al*., 2017). However, besides these formally acknowledged and managed UGS, a multitude of so-called informal urban green spaces (IGS) such as “vacant” lots, street or railway sidings, utility easements, corridors between buildings and riverbanks are typically deprioritized and often represent an underrated piece of the urban nature and urban planning puzzle (Rupprecht & Byrne, 2014). Among these neglected IGS, urban wastelands (or “brownfield” lands) come in all sizes and shapes, and unlike the coherently managed urban parks, they often meet the diverse “nature needs” of their users who, contrary to urban planners, do not view IGS as being “vacant” or as an “empty space” that should be developed (Rupprecht *et al*., 2015; Botzat *et al*., 2016). In their meta-analysis spanning across 37 independent studies, Bonthoux *et al*. (2014) show that wastelands are indeed an essential component of urban biodiversity, particularly for birds (see also Villaseñor *et al*., 2020) and plants (e.g., Godefroid *et al*., 2007), but also for beetles (Coleoptera) (Small *et al*., 2002; Small *et al*., 2003). To date, the explicit contribution of wastelands to the diversity of other groups of organisms relevant to urban ecosystem services provision, such as wild bees (Hymenoptera, Apoidea), remains relatively poorly understood (but see Fischer *et al*., 2016; Twerd & Banaszak-Cibicka, 2019).

In this study, we assess the contribution of the Friche Josaphat, the largest urban wasteland in Brussels, to the diversity of wild bees at the scale of the Brussels-Capital Region. We compiled five years of field surveys to characterize the fauna of this site, and we compare it to the regional checklist of wild bees. Specifically, we use taxonomic, traits-based functional and phylogenetic diversity metrics, as well as null models of community assembly to test if the wild bee species assemblage recorded at our study site is functionally and phylogenetically clustered or over-dispersed (i.e., with significantly less or more similarities among co-occurring species than in a random community, respectively). The importance of environmental filtering as a driver structuring the wild bee community of the Friche Josaphat is discussed, as well as the implications of our results for the conservation of urban bees and urban biodiversity.

## Materials and methods

### Study site

The 24-hectare Friche Josaphat (Figure 1) is currently one of the few remaining wastelands in the Brussels-Capital Region (Belgium; N50.863224, E4.395417), and by far the largest in size. This site is a former railway marshalling yard extending across the Schaerbeek and Evere municipalities; in other words, it is a post-industrial urban fallow now turned into a semi-natural meadow, and one of the largest unfragmented green spaces entirely enclaved in the dense urban matrix of Brussels (Figure 1). After the closure of the Schaerbeek-Josaphat marshalling yard along the Railway Line 26 (Mechelen-Etterbeek-Hal) in 1994, the railway infrastructure was dismantled and the site was subsequently cleaned up, levelled with soil and sand, and turned into a semi-natural grassland in 2013 (Figure 1).

**Figure 1.**
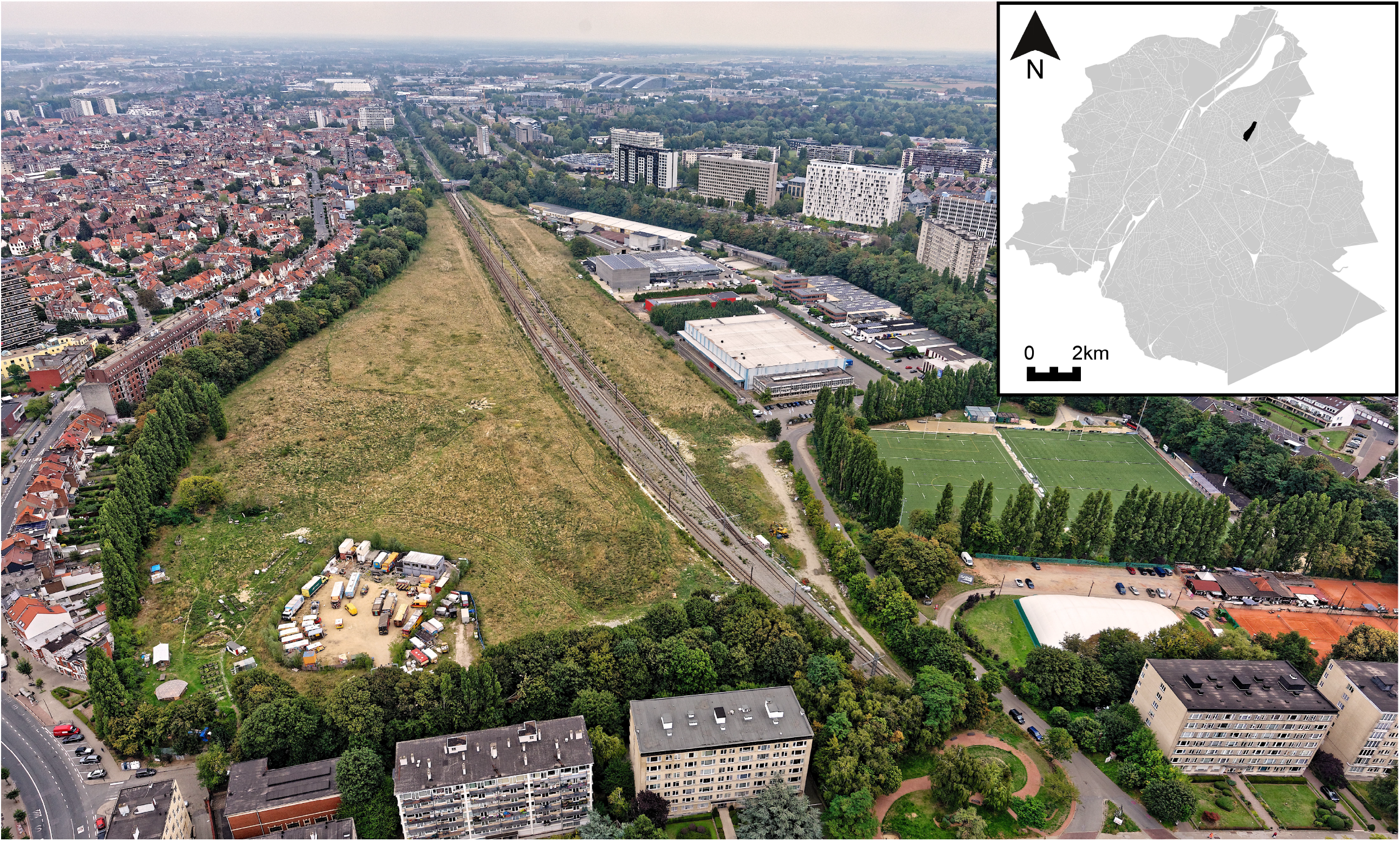
Aerial view of the Friche Josaphat wasteland in Brussels (Belgium) surrounded by a row of tall trees and the railway tracks (Photo © S. Schmitt/Global View - Photographie aérienne), and its location in the Brussels-Capital Region (in the top right corner).

The site is currently owned by the Urban Development Corporation of the Brussels-Capital Region (Société d’Aménagement Urbain, SAU) and according to present-day development plans, the semi-natural meadow will be largely destroyed and turned into impervious surfaces, perhaps at the exception of seven hectares converted into biodiversity-focused embankment (3.0ha), an active linear park (2.8ha) and a relaxation space (1.1ha). This public announcement has stimulated collaborative research among citizens, researchers and local non-profit organisations to document the wildlife conservation value of the Friche Josaphat for a variety of taxonomic groups, with the overarching goal to propose alternative, biodiversity-inclusive and participatory management approaches for the Friche Josaphat site. Our contribution to this collective endeavour was to conduct new field surveys and collect all available and verified records relevant to the wild bees of the Brussels-Capital Region.

### Data collection

Our dataset encompasses abundance records of wild bees obtained between 10.vi.2015 and 31.x.2020 through opportunistic observations, as well as through standardized, targeted biological surveys using a combination of pan traps and insect netting (transect walks) (Westphal *et al*., 2008; Normandin *et al*., 2017; Leclercq *et al*., unpublished; Weekers *et al*., unpublished). The methodology on the use of pan traps for bee surveys is detailed in Vereecken *et al*. (2021).

Opportunistic surveys at the Friche Josaphat by amateur naturalists started in 2015 up to the present day; we compiled all validated observations available through the citizen science platforms Observations.be/Waarnemingen.be (2021), including the date, time, geographic coordinates, field notes, as well as photographs as supporting evidence. Individual records obtained through citizen science surveys relate to a single species, yet they can include the number of specimens of the species observed locally which can amount to several hundreds in the case of a nesting aggregation.

Standardized field surveys consisted in combining insect netting and pan traps (Westphal *et al*., 2008; Leclercq *et al*., unpublished). All bees were then pinned and labelled, then identified down to the species level. Individual records here correspond to pinned specimens.

### Statistical analyses

We first prepared a species accumulation curve by randomly assigning the order of specimens observed (Gotellli & Colwell, 2001) and the *specaccum* function in the ‘‘vegan’’ package (Oksanen *et al*., 2020) to visually assess the adequacy of our wild bee field surveys. This and all following analyses were conducted in RStudio (2020) for R (R Core Team, 2020). We then calculated the total expected species richness (or the number of unobserved species) using a bootstrapping procedure with n=999 random reorganizations of sampling order. Total expected species richness was assessed using Chao (1984), Jack1 (First order jackknife), and Jack2 (Second order jackknife) estimators with the *alpha*.*estimate* function in the “BAT” package (version 2.5.0.) (Cardoso *et al*., 2015) (see Normandin *et al*., 2017 for details).

For functional community structure approaches, we used the methodology described in Vereecken *et al*. (2021): the taxonomic classification and functional traits of wild bee species in the Brussels-Capital Region used in this study are available in Table S1. The mixed matrix of qualitative and quantitative functional traits (between the columns “ITD” and “Diet.breadth”) was converted into a Gower distance matrix with the *gowdis* function in the “FD” package (version 1.0–12) (Laliberté & Legendre 2010; Laliberté *et al*., 2015). We then used the *pcoa* function from the “ape” package (version 5.0) (Paradis & Schliep, 2019) to perform a principal coordinates analysis (PCoA) based on the Gower distance matrix above, and we used the first two principal coordinates to plot the functional space of the Friche Josaphat and the Brussels-Capital Region wild bee communities as convex hulls, following the framework described by Mouillot *et al*. (2013). We excluded species in the subgenus *Micrandrena* (*Andrena*, Andrenidae) from the analyses as they are notoriously challenging to identify and still await a proper revision, and we also excluded *Hylaeus paulus* (Colletidae) (1 specimen collected at the Friche Josaphat) and *Nomada pleurosticta* (Apidae) (1 specimen collected in Brussels outside the Friche Josaphat) because we failed to compute their inter-tegular distance (ITD). We used the *multidimFD* function by Mouillot *et al*. (2013) to characterize the functional β-diversity between the wild bee community of the Friche Josaphat and that of the Brussels-Capital Region by computing the proportion of the nested, multi-dimensional convex hull of the Friche Josaphat (*FRic*, in %, as functional richness or the proportion of functional space filled by species present in the assemblage).

To compare the phylogenetic structure of the Friche Josaphat community to that of the Brussels-Capital Region, we adopted the approach described in Vereecken *et al*. (2021) by building a polytomous, ultrametric tree based on the Linnaean taxonomic hierarchy of wild bees, and we used the “ggtree” package (version 3.12) (Yu *et al*., 2017; Yu, 2020) to visualize the resulting phylogenetic tree with its associated location data.

Last, for both the traits-based functional and phylogenetic approaches, we have computed the Mean (Functional or Phylogenetic) Distance (M(F/P)D), an average for the pairwise (functional or phylogenetic) distances values across all pairs of taxa in a community (in the functional space or across the phylogeny). We also computed a traits-based functional and phylogenetic Mean Nearest Taxon Distance (MNTD), a metric that provides an average of the (functional or phylogenetic) distances between each species and its nearest (functional or phylogenetic) neighbor in the community (Webb *et al*., 2002; see also Dorchin *et al*. 2018). Specifically, we computed the Standardized Effect Sizes (SES; Gotelli & Cabe, 2002) to compare the functional and phylogenetic scores for M(F/P)D and MNTD obtained from the observed community with a randomized null community (n=999). These variables were calculated with the “picante” package (Kembel *et al*., 2010).

## Results and discussion

### The wild bee fauna of the Friche Josaphat in Brussels

Our dataset for the Friche Josaphat comprises 2,507 individual records, representing 7,188 specimens and 127 species of wild bees, as well as the honey bee (*Apis mellifera*) (Table S1). The highest estimation of species richness for the Friche Josaphat was associated with the second order Jackknife estimator (168.84 species), while the lowest was the first order Jackknife estimator (150 species). The Chao estimator indicated the probable presence of 153.67 species at our study site. Collectively, these results, along with the shape of the species accumulation curve reaching a plateau (Figure 2), indicate that we have observed 75.81-85.33% of the estimated species richness at the Friche Josaphat. In terms of taxonomic diversity, the wild bees recorded at the Friche Josaphat belong to six families and 26 genera (Figure 2), and they account for 60.6% of the 210 species recorded in the Brussels-Capital Region between February 1999 and March 2020, or 34.5% of the 345 species assessed recently in the Belgian Red List of Bees (Drossart *et al*., 2019).

**Figure 2.**
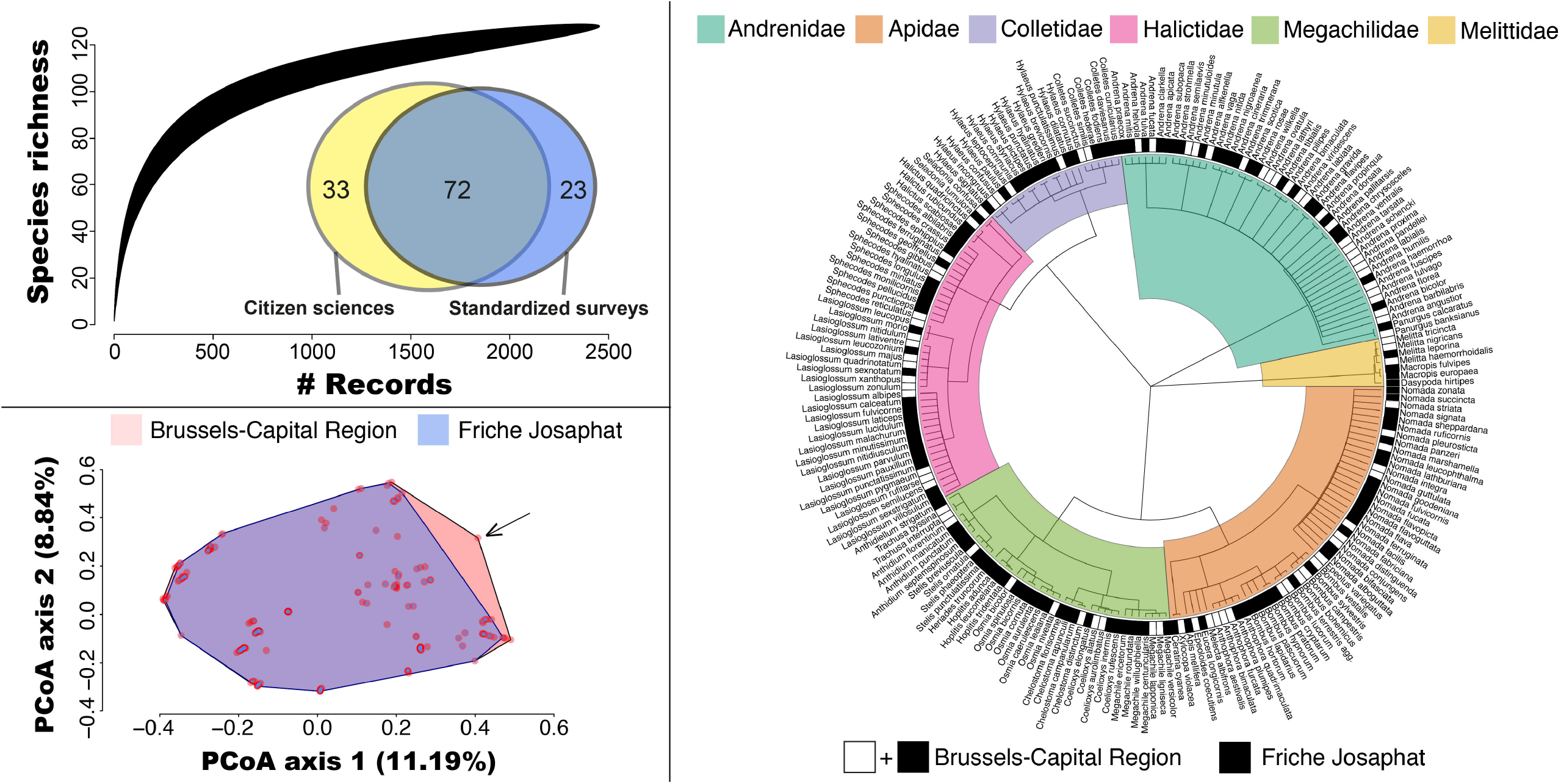
Analysis of the wild bee community structure associated with the Friche Josaphat and the Brussels-Capital Region. ***Top left*:** Species accumulation curve using a bootstrapping procedure with n=999 random reorganizations of sampling order. The mean species accumulation curve of the Friche Josaphat reaches a plateau, and estimators indicate that we have observed 75.81-85.33% of the estimated local species richness, which confirms the adequacy of our field surveys. The Venn diagram indicates the number of unique and shared species recorded by citizen science and standardized surveys. ***Bottom left*:** The pink convex hull represents 100% of the multi-dimensional functional space occupied by all species recorded in the Brussels-Capital Region, and the blue convex hull represents 92.21% of the multi-dimensional functional space occupied by species from the Friche Josaphat. Pink and blue circles are species of wild bees associated with each community; the arrow indicates the position of *Coelioxys aurolimbata* (Hym. Megachilidae), a uniquely large cuckoo bee species associated with a host (*Megachile ericetorum* (Hym. Megachilidae) displaying strong preference (i.e., oligolecty) for host plants in the family *Fabaceae*. All other *Coelioxys* species found in the Brussels-Capital Region or in the Friche Josaphat are associated with pollen generalist (i.e., polylectic) hosts. ***Right***: Phylogenetic classification of wild bees belonging to the six families recorded in the Brussels-Capital Region (black and white squares) and in the Friche Josaphat (black squares only).

The 12 most common species are illustrated and listed in Figure S1 along with their abundance in the dataset; they represented 74.9% of all samples recorded. Our records encompass seven wild bee species of conservation concern at the scale of Belgium: these include the nationally “Vulnerable” species *Eucera longicornis* (Apidae) and the “Near threatened” species *Andrena bimaculata* (Andrenidae), *Bombus hortorum* (Apidae), *Coelioxys rufescens* (Megachilidae), *Osmia aurulenta* (Megachilidae), *Osmia spinulosa* (Megachilidae) and *Stelis phaeoptera* (Megachilidae) (Drossart *et al*., 2019). The Friche Josaphat is also home to species threatened with extinction at the European scale, such as the “Vulnerable” *Colletes fodiens* (Colletidae) and the “Near threatened” *Lasioglossum sexnotatum* (Halictidae) (Nieto *et al*., 2014) (Table S1). We also noted the presence of four species recorded in the Brussels-Capital Region that are only known from the Friche Josaphat so far, namely *Hylaeus paulus* (Colletidae), *Anthidium punctatum* (Megachilidae) and *Osmia aurulenta* (Megachilidae). Moreover, the record of *Anthidium septemspinosum* (Megachilidae) through citizen sciences surveys was not only unique to the Friche Josaphat, but also a new addition to the Belgian checklist of wild bees (Vereecken *et al*., unpublished).

A total of 72 bee species were recorded both through citizen sciences and standardised field surveys at the Friche Josaphat. The citizen sciences records included another 33 additional wild bee species not detected through standardized field surveys, whereas the standardised field surveys helped adding 23 species not detected through the citizen sciences surveys (Figure 2).

We hypothesize that the high biodiversity of wild bees highlighted at the Friche Josaphat stems from several important factors, including (i) the biodiversity-friendly management of the Friche Josaphat since 2013, (ii) the strong complementarity of citizen science and standardised methods in biodiversity surveys, particularly when they aim at maximizing the number of species recorded in a check-list format, and (iii) the proximity of the railway and railway edges to the Friche Josaphat (Figure 1). Indeed, Linear Transport Infrastructures such as railways, but also highways, waterways and power transmission lines are increasingly acknowledged as important biodiversity corridors for invertebrates, including for pollinators and wild bees in particular (e.g., Wojcik & Buchmann, 2012; Wagner *et al*., 2014; Hill & Bartomeus, 2016; Steinert *et al*., 2020).

Our results on the traits-based functional community structure illustrate that the Friche Josaphat represents 92.21% of the functional space occupied by all wild bee species recorded in the Brussels-Capital Region (Figure 2 & Figure S2). Our analysis of the community structure of bees indicates that, compared to the Brussels-Capital Region, the diversity of the Friche Josaphat community is significantly reduced taxonomically (211 vs. 128 species), functionally (FD observed = 19.847 vs. 14.304; *p*-value=0.03) and phylogenetically (PD observed = 10.365 vs. 8.759; *p*-value=0.0280) (see also Figure 2). However, we found no significant difference between the Brussels-Capital Region and the Friche Josaphat communities when we computed the Mean (Functional or Phylogenetic) Distance (M(F/P)D) (MFunctD z-score=-0.027 and *p*-value=0.461; MPhyloD z-score=-1.536 and *p*-value=0.077), or the traits-based functional and phylogenetic Mean Nearest Taxon Distance (MNTD) (MNTD_traits_ z-score=-1.130 and *p*-value=0.124; MNTD_phylo_ z-score=-1.228 and *p*-value=0.115). These results illustrate that the Friche Josaphat encompasses a randomly nested subset of the wild bee fauna of the Brussels-Capital Region from a functional and phylogenetic perspective, and also suggest a negligible role of environmental filtering towards certain combinations of traits or taxonomic groups in the community assembly process. As such, the Friche Josaphat is therefore currently the richest semi-natural site at the regional level, and the fact that its functional and phylogenetic structure is not significantly different from random communities makes it an ideal site to “showcase” the diversity of urban bees in Brussels.

To date, the most biodiverse site in the Brussels-Capital Region for wild bees was the flower-rich, 5.3 ha “Jean Massart” botanical garden, a Natura 2000 site at Auderghem, which is home to 112 species (Pauly, 2019; surveyed between 1975-2016). Other formally recognized UGS relevant for wild bee diversity include several nature reserves such as the Vogelzangbeek (Anderlecht, 20 ha, 51 spp.), the Scheutbos (Molenbeek, 66 ha, 80 spp.) or the Moeraske (Evere, 14 ha, 69 spp.) according to the citizen science online platforms Observations.be/Waarnemingen.be. The Friche Josaphat also turns out to be one of Belgium’s most diverse sites for wild bees, since the only other known “hotspots” at the scale of Belgium include sites with comparable species counts, but that are much larger nature reserves and/or sites that have been surveyed for several decades (e.g. the Belvédère and Fond-Saint-Martin nature reserve at Han-sur-Lesse with 131 spp. surveyed between 1951-2017 (Pauly & Vereecken, 2019), or the 500+ ha nature reserves Most (a peaty depression) and Keiheuvel (a land dune area) at Balen, with 136 spp. surveyed between 2012-2017 (M. Jacobs, unpublished report).

### The future of urban wastelands and other informal urban green spaces

Collectively, our results have allowed us to confirm recent findings by Twerd & Banaszak-Cibicka (2019) that urban wastelands represent a hitherto underrated and largely overlooked category of UGS with a high potential for the conservation of wild bees in metropolises. More coordination is required among stakeholders to identify the local factors contributing to the high biodiversity of wild bees at the Friche Josaphat: for example, Strauss & Biedermann (2006) have provided evidence that phytophagous insects in biodiverse urban brownfields have clear preferences for certain successional stages of the vegetation. The conservation of a rich local species pool within a city therefore requires coordinated and evidence-based management of the vegetation, and perhaps the maintenance of a mosaic of (all) successional vegetation stages that provide the key host plants of ecologically specialized and generalized species alike. In the case of the Friche Josaphat, addressing these challenges requires an integrated landscape approach tailored to the ecological requirements of the targeted species (see Table S1) (Wilson & Jamieson, 2019) within the ecological network of important UGS (Ayers & Rehan, 2021) such as the neighbouring cemetery of Brussels and the Josaphat Park. Another category of IGS or “vacant land” that received increasing attention over the past decade are urban agriculture plots such as community gardens. Results from recent studies indicate that they too have the potential to harbour particularly high levels of species richness for wild bees and other components of wildlife in urban centres (Normandin *et al*., 2017; Turo *et al*., 2021; Vereecken *et al*., 2021).

“Formal” or “conventional” UGS such as parks are (increasingly) expensive to maintain, and they often fail to satisfy the urban dwellers’ diverse needs. In a context of ever-increasing urbanisation pressure, where spatial (conservation) prioritization using appropriately chosen objectives is a pressing priority, biodiverse and highly multi-functional IGS should therefore be urgently provided with a formal status within the mosaic of UGS. Indeed, their uncertain legal, socio-economic, and ecological status represent major obstacles in realizing these IGS’ full societal and environmental potential (e.g., Rupprecht & Byrne, 2014) and severely limit our capacity to develop wildlife-inclusive urban designs (Apfelbeck *et al*., 2020). Envisioning participatory management approaches of IGS for urban environmental planning and recreation is of pivotal importance if we are to safeguard their natural capital and the multiple ecosystem services they provide, including for the physical and mental well-being of urban citizens.

## Acknowledgements

We are grateful to T. Gaudaire, T. Titeux, D. Burhin, H. Simon, B. Pasau, W. Vertommen and W. Proesmans for their dedicated contribution to data collection in the field, to L. Motquin and B. De Boeck for their support, and to the Société d’Aménagement Urbain (SAU) for allowing access to the Friche Josaphat during our biodiversity surveys. This study was made possible with financial support from Bruxelles Environnement (BE/IBGE) in the framework of the “Atlas of the Wild Bees of Brussels” (WildBnB.brussels 2019-2021), as well as from the FNRS/FWO joint programme “EOS — Excellence Of Science” for the project “CliPS: Climate change and its impact on Pollination Services (project 30947854)” to NJV, TW, NL and LM. NJV supervised the sampling, led the writing of the manuscript as well as the statistical analyses; all authors contributed to the collection, analyses and interpretation of data, and reviewed, edited and approved the final version of the manuscript prior to submission.

