## Supplementary figures for "Five years of citizen science and standardized field surveys reveal a threatened urban Eden for wild bees in Brussels, Belgium"

**SUPPORTING INFORMATION**

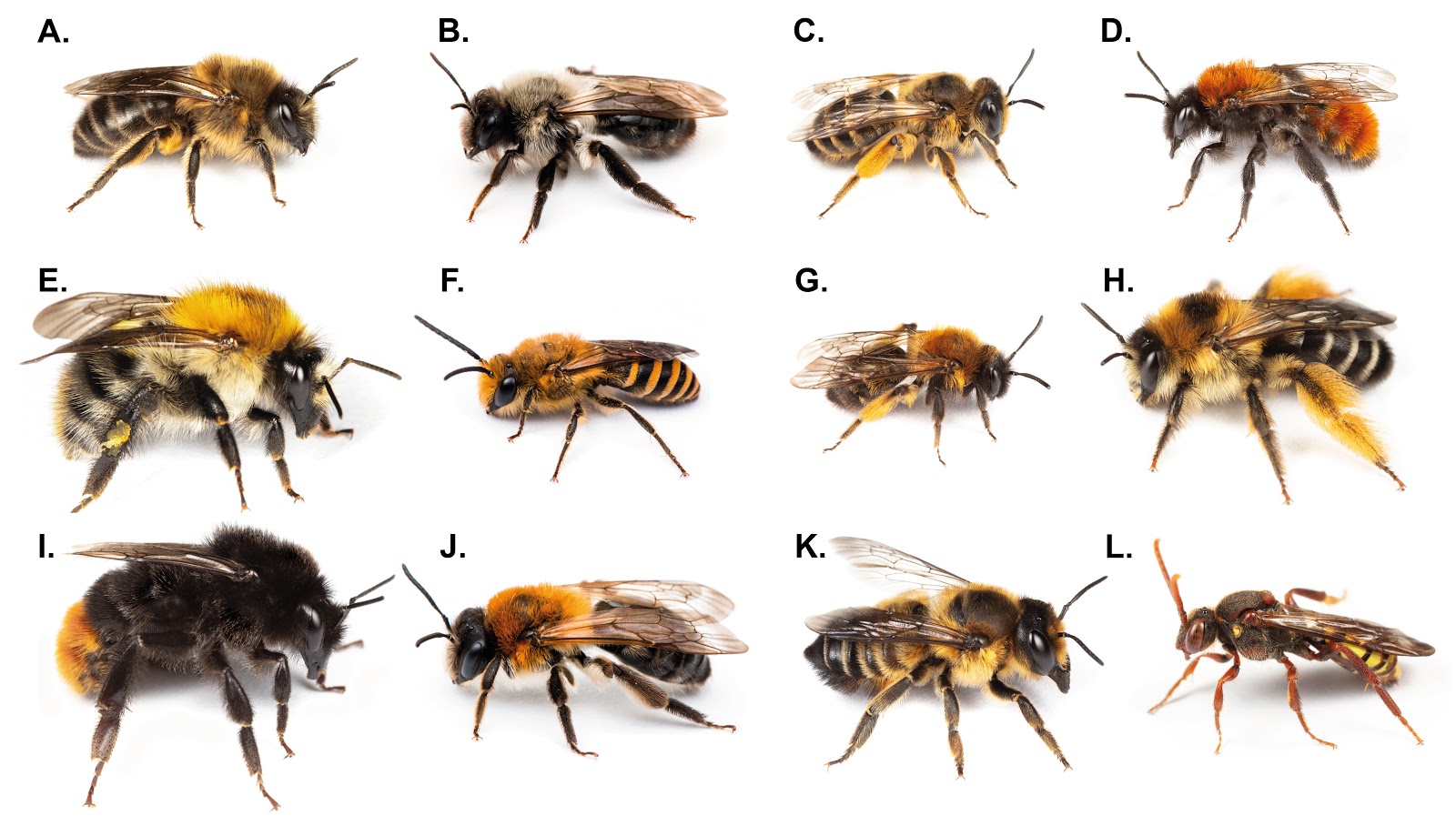

**Figure S1.** Top 12 of the most frequently recorded wild bee species in our dataset for the Friche Josaphat, collectively making up 74.9% of our 7,188 records. **A.** *Colletes cunicularius* (Colletidae) (n=2,230 specimens); **B.** *Andrena vaga* (Andrenidae) (n=1,172 specimens); **C.** *Andrena flavipes* (Andrenidae) (n=526 specimens); **D.** *Andrena fulva* (Andrenidae) (n= 311 specimens); **E.** *Bombus pascuorum* (Apidae) (n= 239 specimens); **F.** *Colletes hederae* (Colletidae) (n=223 specimens); **G.** *Andrena bicolor* (Andrenidae) (n=152 specimens); **H.** *Dasypoda hirtipes* (Melittidae) (n=144 specimens); **I.** *Bombus lapidarius* (Apidae) (n=112 specimens); **J.** *Andrena nitida* (Andrenidae) (n=89 specimens); **K.** *Megachile willughbiella* (Megachilidae) (n=89 specimens); **L.** *Nomada signata* (Apidae) (n=87 specimens). All photographs ©N.J. Vereecken, except **B.-F.-J.** ©S. De Greef.

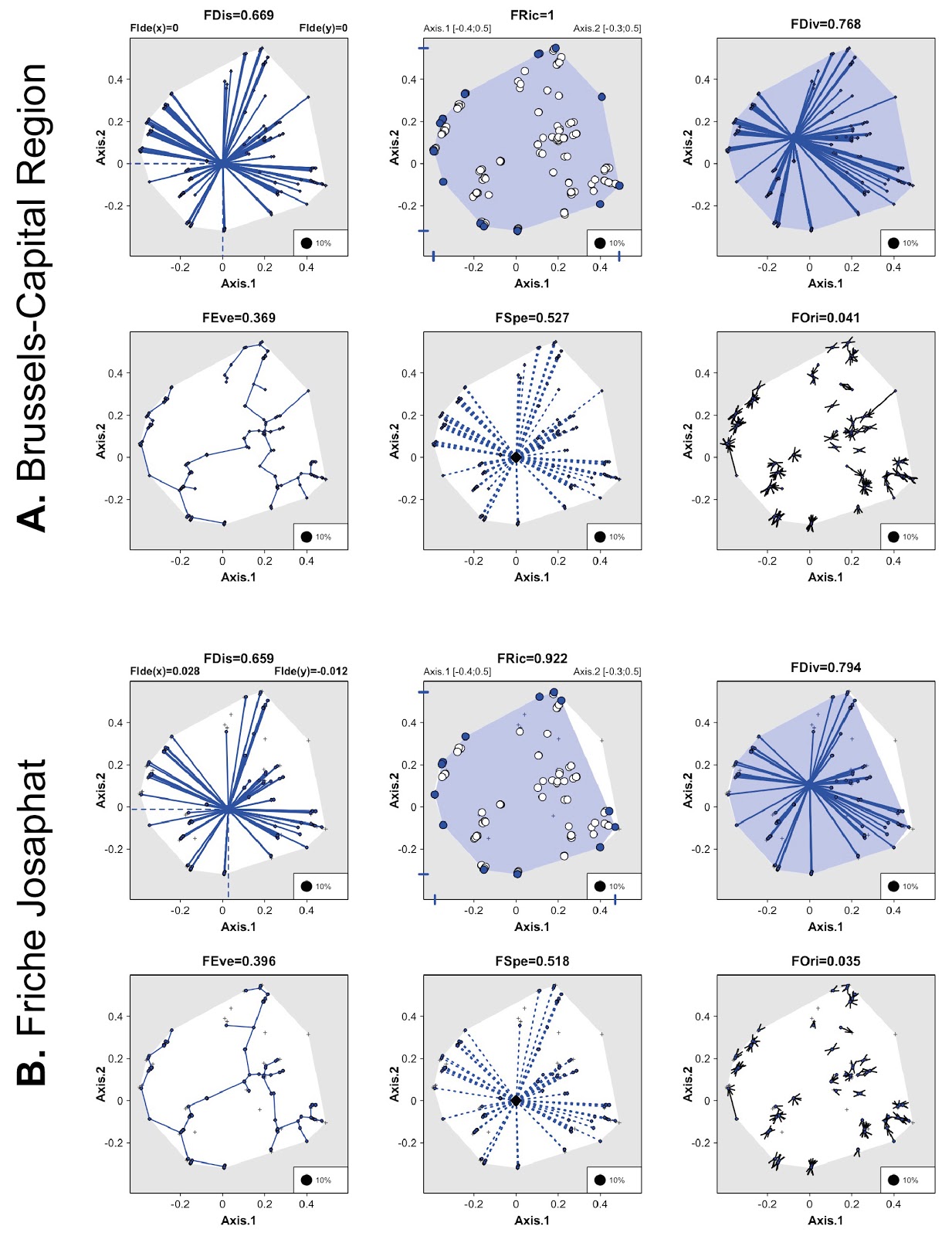

**Figure S2.** Functional community structure similarity between **A.** the Brussels-Capital Region (upper six figures) and **B.** the Friche Josaphat (lower six figures) as illustrated by metrics computed follow Mouillot *et al.* (2013) with a presence-absence matrix only, and including functional dispersion (*FDis*, i.e., changes in the deviation of species trait values from the center of the functional space filled by the community (i.e., the mean distance to the mean trait values of the community), functional richness (*FRic*, i.e., the portion of the functional space filled by species communities), functional divergence (*FDiv*, i.e., the proportion of the total number of species supported by the species with the most extreme functional traits), functional evenness (*FEve*, i.e., a measure of the modifications in the regularity of species distributions in the functional space (along the shortest minimum spanning tree linking all the species), functional specialization (*FSpe*, i.e., illustrating the patterns of generalist species (i.e., species close to the center of the functional space, here linking all species) or specialist species (i.e., having extreme trait combinations), and functional originality (*FOri*, i.e., how changes in the patterns of co-occurring species modify the functional redundancy between species (i.e., black lines are minimal functional distances among species pairs). The white shape in the lower panel represents 100% of the functional space occupied by all species recorded in the Brussels-Capital Region, and the blue shape represents the amount of functional space occupied by species from the Friche Josaphat. White and blue circles are species assigned to vertices in the multidimensional space. Small black crosses represent the position of all species within the functional space.

**Supplementary Table 1**.

The file “SI Table 1.xlsx” contains a list of behavioural and ecological traits of wild bee species, along with their Linnaean classification, covering the fauna of the Brussels-Capital Region and specifying (in the column “Pres.Friche”) if, in the framework of this study, the species were recorded only from the Friche Josaphat, or from the Brussels-Capital Region. The dataset also includes a column listing the IUCN status of each species according to the IUCN Red List of bees for Europe (in the column “EU.IUCN.Status”, following Nieto *et al.* 2014) and for Belgium (in the column “BE.IUCN.Status”, following Drossart *et al.* 2019). All other traits used are described in Vereecken *et al.* (2021). The inter-tegular distance (ITD) of *Hylaeus paulus* (Colletidae) and *Nomada pleurosticta* (Apidae) were unavailable at the time of the study, and they are therefore marked as “NA” in the dataset.
